# Proximity-dependent proteomics reveals extensive interactions of Protocadherin-19 with regulators of Rho GTPases and the microtubule cytoskeleton

**DOI:** 10.1101/2020.09.11.293589

**Authors:** Michelle R. Emond, Sayantanee Biswas, Matthew L. Morrow, James D. Jontes

## Abstract

Protocadherin-19 belongs to the cadherin family of cell surface receptors and has been shown to play essential roles in the development of the vertebrate nervous system. Mutations in human Protocadherin-19 (*PCDH19*) lead to *PCDH19* Female-limited epilepsy (*PCDH19* FLE) in humans, characterized by the early onset of epileptic seizures in children and a range of cognitive and behavioral problems in adults. Despite being considered the second most prevalent gene in epilepsy, very little is known about the intercellular pathways in which it participates. In order to characterize the protein complexes within which Pcdh19 functions, we generated Pcdh19-BioID fusion proteins and utilized proximity-dependent biotinylation to identify neighboring proteins. Proteomic identification and analysis revealed that the Pcdh19 interactome is enriched in proteins that regulate Rho family GTPases, microtubule binding proteins and proteins that regulate cell divisions. We cloned the centrosomal protein Nedd1 and the RacGEF Dock7 and verified their interactions with Pcdh19 *in vitro*. Our findings provide the first comprehensive insights into the interactome of Pcdh19, and provide a platform for future investigations into the cellular and molecular biology of this protein critical to the proper development of the nervous system.

## INTRODUCTION

The cadherin superfamily comprises a diverse array of cell surface glycoproteins defined by the presence of two or more repeats of a ∼110 amino acid cadherin domain (Hirano and Takeichi, 2012; Hulpiau and van Roy, 2011). The protocadherins are the largest subgroup within the cadherin superfamily and can be subdivided into the clustered-protocadherins (c-Pcdhs) and the non-clustered protocadherins (nc-Pcdhs). The nc-Pcdhs consist largely of the δ1-pcdh (*pcdh1, pcdh7, pcdh9* and *pcdh11*) and δ2-pcdh (*pcdh8, pcdh10, pcdh17, pcdh18* and *pcdh19*) subfamilies, as well as a small number of orphan protocadherins, such as *pcdh12, pcdh20* and *pcdh24* (Hulpiau and van Roy, 2011). Mutations in the δ-pcdhs, both δ1- and δ2-pcdhs, are involved in human disease, with particular relevance to neurodevelopmental disorders and cancer (Berx and van Roy, 2009; Hirano and Takeichi, 2012; Redies et al., 2012; van Roy, 2014). For example, epilepsy is associated with mutations in both *PCDH7* (Lal et al., 2015) and *PCDH19* (Depienne et al., 2009; Dibbens et al., 2008; Lal et al., 2015), while *PCDH8, PCDH10* and *PCDH19* are implicated in autism spectrum disorders (Butler et al., 2015; Morrow et al., 2008; Piton et al., 2011). In addition, a deletion of *PCDH18* in one patient was associated with severe developmental delay and cognitive disability (Kasnauskiene et al., 2012). Finally, mutations in *PCDH12* are associated with a congenital microcephaly (Aran et al., 2016; Vineeth et al., 2019).

Several lines of evidence additionally reveal a strong involvement of the δ-pcdhs in cancer, with several of these genes implicated as tumor suppressors (Berx and van Roy, 2009; van Roy, 2014). Numerous studies have shown the suppression of δ-pcdh expression by promoter methylation, both in carcinomas and in cancer-derived cell lines (Tang et al., 2012; Yin et al., 2016; Zhang et al., 2017). In several cases, the loss of δ-pcdh expression correlates with poor prognosis (Bing et al., 2018; Cao et al., 2018; Dang et al., 2016; Lin et al., 2018). In addition, evidence for a tumor suppressive role of the δ-pcdhs has been obtained *in vitro*, as their forced expression limits tumor growth (Xu et al., 2015; Yin et al., 2016; Zong et al., 2017). Supporting a role for δ-pcdhs as tumor suppressors, recent studies suggest that the δ-pcdhs can regulate cell proliferation. Both Pcdh11 and Pcdh19 have been shown to be expressed in neural progenitor cells, and loss of these proteins results in increased proliferation and neurogenesis (Cooper et al., 2015; Zhang et al., 2014). Similarly, *pcdh19* is a target of miR-384, and increased copies of miR-384 result in increased neurogenesis (Fujitani et al., 2016). This increase can be rescued by forced expression of Pcdh19.

Despite the clear importance of the δ-pcdhs to both development and disease, relatively little is understood of their molecular function. Like the classical cadherins, the protocadherins can mediate homophilic cell adhesion (Bisogni et al., 2018; Cooper et al., 2016; Harrison et al., 2020). However, in contrast to the classical cadherins, the protocadherins do not have conserved binding sites for the armadillo proteins p120-catenin or for β-catenin, which links cadherins to the cytoskeleton though α-catenin (Ozawa et al., 1989; Ozawa and Kemler, 1992). A relatively small number of cytoplasmic effectors have been reported. The δ1-pcdh Pcdh7/NF-pcdh interacts with the protein TAF1/Set in the contexts of both neural tube closure and axon outgrowth (Heggem and Bradley, 2003; Piper et al., 2008). The δ1-pcdhs have also been shown to interact with protein phosphatase 1α (Vanhalst et al., 2005; Yoshida et al., 1999). In hippocampal neurons, the δ2-pcdh Pcdh8/Arcadlin interacts with the kinase TAO2β to regulate internalization of N-cadherin at the synapse (Yasuda et al., 2007). Most recently, the δ2-pcdhs, as well as Pcdh9, were found to interact with the WAVE complex through a conserved WIRS site (WAVE Interacting Receptor Site) (Chen et al., 2014). This interaction appears to be important for contact-dependent cell motility and axon growth (Biswas et al., 2014; Hayashi et al., 2014; Nakao et al., 2008; Tai et al., 2010).

To have a better understanding of δ-protocadherin function and the full range of their cellular roles, it is necessary to have a more complete view of their downstream intracellular networks. To address this issue, we used proximity-dependent biotinylation and proteomics analysis to determine the proxeome of Pcdh19 fused to the promiscuous biotin ligases, BioID (Roux et al., 2012) or BioID2 (Kim et al., 2016). In addition to recovering interactions with previously identified binding partners, such as members of the WAVE complex, we identified strong interactions with the actin cytoskeleton, regulators of Rho GTPases, and the microtubule cytoskeleton and regulators of cell division. The identification of this downstream network is consistent with a role for the δ-pcdhs in the regulation of cell proliferation and provides new insights into the biological function of the δ-protocadherin family.

## RESULTS

### Identification of the Pcdh19 interactome using BioID or BioID2

In order to identify proteins that interact with zebrafish Pcdh19, we performed proximity-dependent biotinylation using fusions to promiscuous biotin ligases (Roux et al., 2012). Proteins that are present within ∼10nm of the biotin ligase will be labeled with biotin on primary amines, allowing them to be isolated on streptavidin-conjugated magnetic beads. We fused BioID (Roux et al., 2012) or BioID2 (Kim et al., 2016) to the carboxy-terminus of the Pcdh19 intracellular domain and expressed the full fusion proteins in HEK293 cells (**Figure 1A,B**). To confirm the biotinylation of Pcdh19 interacting proteins, we performed SDS-PAGE and Western blotting on cell lysates. HEK293 cells were transfected with Pcdh19-BioID2-HA in the presence or absence of added biotin in cell culture media. Cell lysates were run on an SDS-PAGE gel and immunoblotted using an antibody against the HA epitope (**Figure 1B**) or added to streptavidin magnetic beads before detection with streptavidin-HRP (**Figure 1C**). Western blotting of the cell lysates indicated that significant biotinylation was evident only in cells that had both biotin added to the media and that were expressing the Pcdh19-BioID2-HA protein. To further characterize protein biotinylation by the fusion protein, we transfected cells with Pcdh19-BioID2-HA and performed immunocytochemistry with an anti-HA antibody and a Streptavidin-HRP to label biotinylated proteins. In the presence of both Pcdh19-BioID2-HA and biotin, cells exhibited robust biotinylation of cytosolic proteins (**Figure 1D-F**). We observed minimal biotinylation in the absence of Pcdh19-BioID2-HA (**Figure 1G-I**).

**Figure 1.**
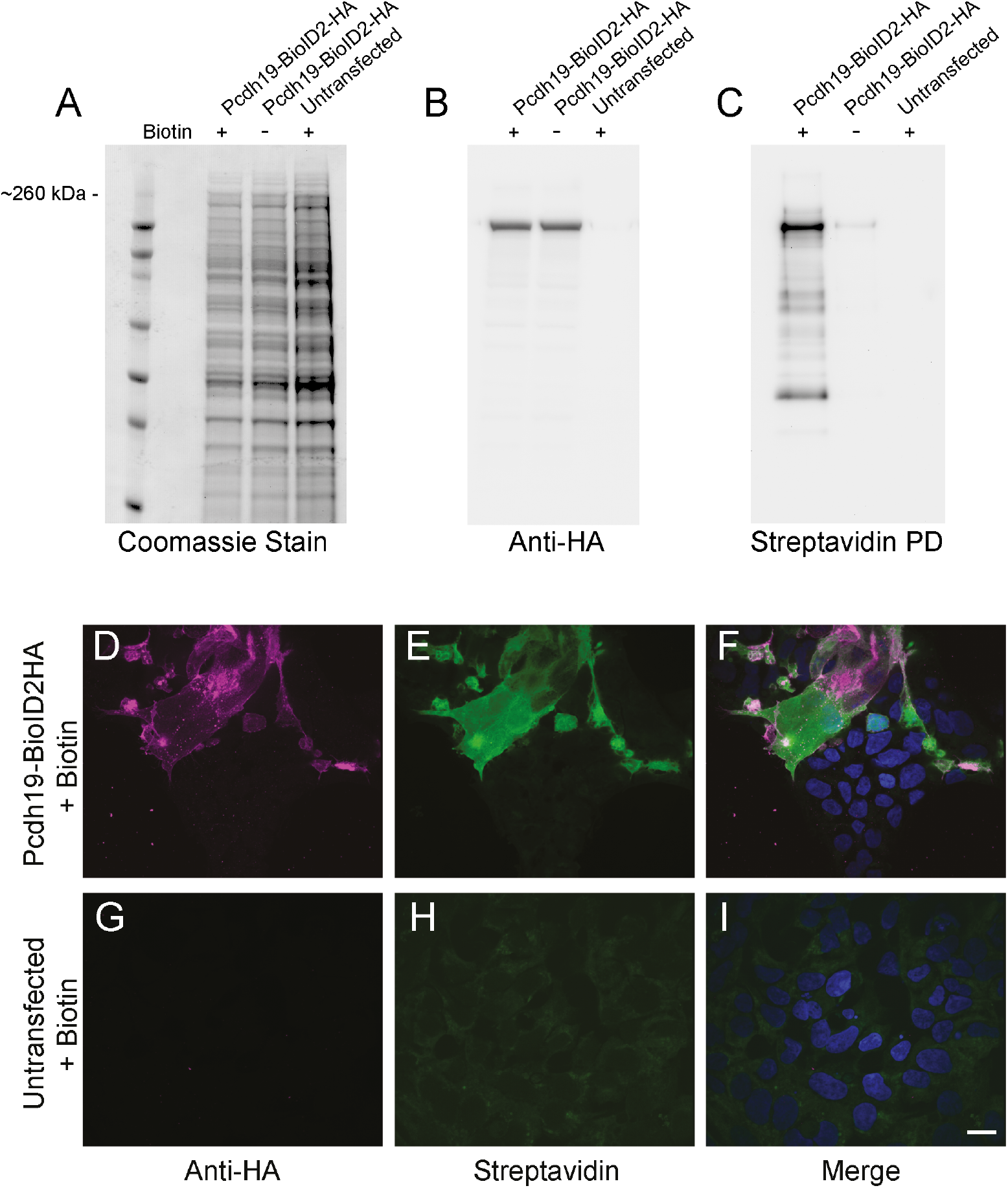
Characterization of Pcdh19-BioID2 fusion protein. **A**. Total cell lysates were separated on an SDS-PAGE gel and stained with Coomassie from untransfected HEK293 cells and cells transfected with Pcdh19-BioID2-HA, both in the presence or absence of added biotin. **B**. Western blot analysis of HEK293 cells transffected with Pcdh19-BioID2-HA in the presence and absence of biotin and stained with anti-HA antibodies. Untransfected cells treated with biotin served as a control. **C**. Biotinylated proteins were pulled down using streptavidin-conjugated magnetic beads and detected using a peroxidase-conjugated streptavidin. Labeled proteins are only detected in the presence of both Pcdh19-BioID2-HA and added biotin. **D-F**. Immunocytochemistry of HEK293 cells transfected with Pcdh19-BioID2-HA. In the presence of added biotin, cells that express Pcdh19-BioID2-HA (**D**) exhibit biotinylation of intracellular proteins, as detected by Streptavidin-peroxidase (**E**,**F**). **G-I**. In the absence Pcdh19-BioID2-HA (**G**), biotin alone does not lead to significant labeling of cellular proteins is detected (**H**,**I**). Scale bar = 10 *μ*m.

To identify the Pcdh19 interactome, we prepared cell extracts from HEK293 cells that had been transfected with either Pcdh19-BioID-HA or Pcdh19-BioID2-HA and grown for 24 hours in the presence of biotin. Biotinylated proteins were bound to Streptavidin-conjugated magnetic beads and washed extensively, before analysis by capillary-liquid chromatography-nanospray tandem mass spectrometry (Capillary-LC/MS/MS). We performed five independent experiments (two with BioID and three with BioID2). From the resulting protein lists (**Supplemental File 1**), we removed common contaminants, including keratins, nuclear proteins and ribosomal proteins. We further filtered the list by comparing our list to a database of common contaminants (www.crapome.org), eliminating any protein that occurred in >15% of comparable proteomics experiments. We then retained proteins that were found in 3 of the 5 experiments, leaving 159 proteins (**Supplemental File 2**). The δ2-protocadherins, which includes Pcdh19, have previously been shown to interact with the WAVE complex (Biswas et al., 2014; Chen et al., 2014; Hayashi et al., 2014; Nakao et al., 2008; Tai et al., 2010), which promotes Rac1-dependent actin assembly (Ismail et al., 2009). The WAVE complex consists of five components, Cytoplasmic FMR1 interacting protein 1 or 2 (CYFIP1/2), Nck associated protein 1 (Nap1), Abl interacting protein 1 or 2 (Abi1/2), Wiskott-Aldrich syndrome family member 1, 2 or 3 (WASF1/2/3) and Hematopoetic stem/progenitor cell protein 300 (HSPC300) (Chen et al., 2014; Ismail et al., 2009). Of these components, Cyfip1/2, Abi2 and WAVE2 (WASF2) were retained in our filtered list, indicating that this approach effectively identifies Pcdh19 interacting proteins.

### The Pcdh19 interactome is enriched in regulators of Rho family GTPases and regulators of the microtubule cytoskeleton

In order to understand the intracellular pathways downstream of Pcdh19, our list of 159 proteins was classified by Gene Ontology analysis (**Figure 2**). As the δ2-pcdhs are known to interact with the WAVE complex, which requires activated Rac1 to promote actin assembly, we anticipated an enrichment of actin associated proteins, as well as regulators of Rho GTPases. Consistent with these expectations, we found that both actin associated proteins and Rho family GTPase binding proteins were enriched among candidate proteins (**Figure 2A**), with cytoskeletal protein binding being the largest cluster of proteins. In addition to the components of the WAVE complex, Cyfip1/2, Abi1/2 and Wasf2, other actin binding proteins included Formin homology 2 domain containing protein 1 (FHOD1), Anilin (ANLN), Shroom3 (SHRM3), and Afadin (AFDN) (**Table 1**). Among regulators of Rho GTPases were Dedicator of cytokinesis protein 7 (DOCK7), Rho guanine exchange factor 2(ARHGEF2), and T-lymphoma and metastasis including protein 1 (TIAM1) (**Table 2**). Other prominent proteins included cadherin and β-catenin binding proteins (**Figure 2A**).

**Table 1.** Actin-associated proteins.

| Gene ID | Protein Name |
| --- | --- |
| ABI2 | Abl Interactor-2 |
| ANLN | Anilin |
| CLASP2 | CLIP-associated protein 2 |
| COBL | Cordon Bleu |
| CYFIP1 | Cytoplasmic FRM1-interacting protein 1 |
| EPB41L5 | Band 4.1-like protein 5 |
| FHOD1 | FH1/FH2 domain-containing protein 1 |
| MACF1 | Microtubule-actin cross-linking factor 1 |
| MAGI1 | Membran-associated guanylate kinase, WW and PDZ domain-containing protein-1 |
| MYO6 | Unconventional myosin-VI |
| SYNE2 | Nesprin-2 |
| SHRM3 | Shroom-3 |
| WASF2 | Wiskott-Aldrich syndrome protein family member 2 |

**Table 2:**
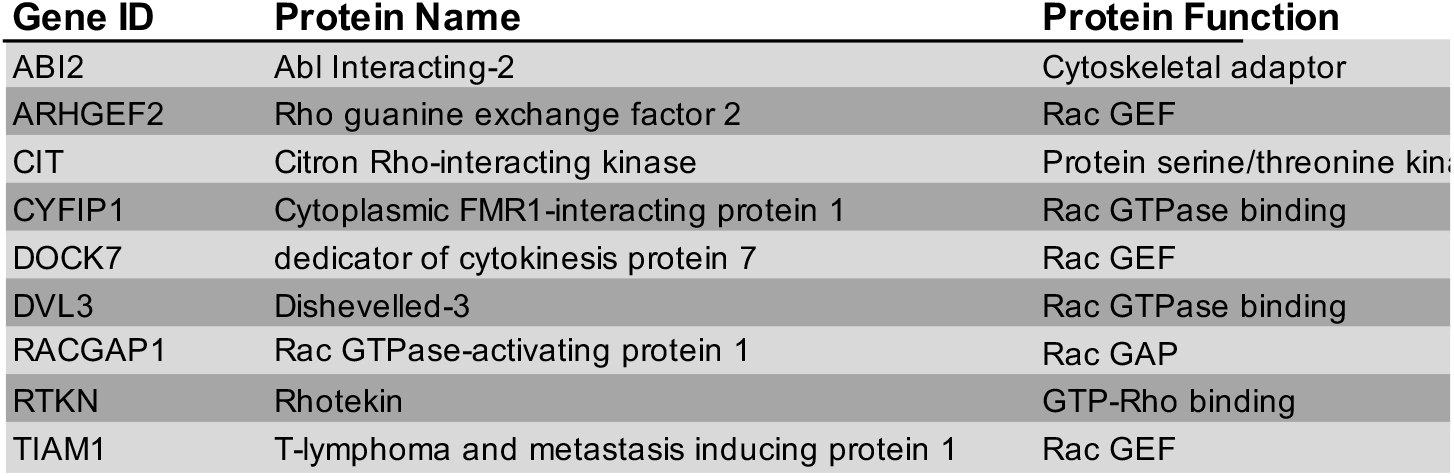
Regulators of Rho GTPases.

| Gene ID | Protein Name | Protein Function |
| --- | --- | --- |
| ABI2 | Abl Interacting-2 | Cytoskeletal adaptor |
| ARHGEF2 | Rho guanine exchange factor 2 | Rac GEF |
| CIT | Citron Rho-interacting kinase | Protein serine/threonine kinase |
| CYFIP1 | Cytoplasmic FMR1-interacting protein 1 | Rac GTPase binding |
| DOCK7 | dedicator of cytokinesis protein 7 | Rac GEF |
| DVL3 | Dishevelled-3 | Rac GTPase binding |
| RACGAP1 | Rac GTPase-activating protein 1 | Rac GAP |
| RTKN | Rhotekin | GTP-Rho binding |
| TIAM1 | T-lymphoma and metastasis inducing protein 1 | Rac GEF |

**Figure 2.**
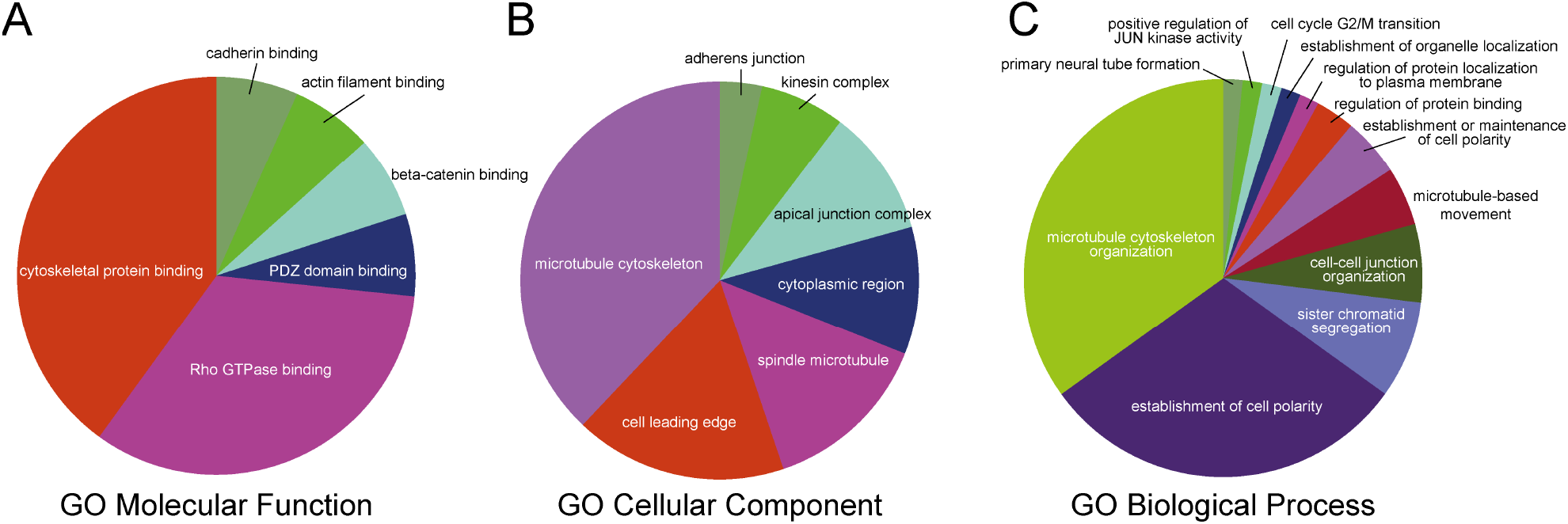
Functional analysis of Pcdh19 interactome. Gene Ontology (GO) analsysis was used to cluster Pcdh19 interacting proteins based on molecular function (**A**), subcellular localization (**B**) or biological process (**C**). **A**. The largest proportions of Pcdh19 interacting proteins were those characterized as cytoskeletal binding proteins and Rho GTPase binding proteins. **B**. Pcdh19 interacting proteins were enriched in proteins that interact with the microtubule cytoskeleton, including spindle microtubules and kinesins. **C**. At the functional level, the Pcdh19 interactome is enriched in proteins associated with microtubules and microtubule-based processes, such as cell polarity, microtubule-based movements, organelle localization and cell cycle regulation.

While many of the identified proteins localize to the leading edge, as would be expected from the linkage of Pcdh19 to actin assembly and cell motility, our list of interactors was also enriched for proteins associated with the microtubule cytoskeleton and spindle (**Figure 2B**; **Table 3**). Neither Pcdh19, nor other δ2-pcdhs, have previously been linked to the microtubule cytoskeleton. Classification of proteins by biological process similarly revealed an enrichment for proteins involved in microtubule organization and the establishment of cell polarity, as well as other microtubule-based processes, such as chromatid segregation, microtubule-based movement and the cell cycle G2/M transition (**Figure 2C**). Overall, while our proteomics results are consistent with a role in contact-dependent actin assembly and cell motility, they also suggest that δ2-pcdhs may also be involved in regulating microtubule cytoskeletal organization.

**Table 3:** Microtubule binding proteins.

| Gene ID | Protein Name |
| --- | --- |
| ARHGEF2 | Rho guanine nucleotide exchange factor 2 |
| CAMSAP1 | Calmodulin-regulated spectrin-associated protein 1 |
| CDK5RAP2 | CDK5 regulatory subunit-associated protein 2 |
| CLASP2 | CLIP-associated protein 2 |
| MACF1 | Microtubule-actin cross-linking factor 1 |
| MAST2 | Microtubule-associated serine/threonine kinase 2 |
| PLK1 | Serine/threonine-protein kinase PLK1 |
| PRC1 | Protein regulator of cytokinesis 1 |
| RACGAP1 | Rac GTPase-activating protein 1 |
| TIAM1 | T-lymphoma invasion and metastasis-inducing protein 1 |

### Verification of selected interacting partners of Pcdh19

Functional analysis of our list of candidate Pcdh19 interacting proteins showed an enrichment in proteins associated with the microtubule network and with regulators of Rho family GTPases. To validate these observations, we selected candidates from each of these groups to verify the putative interaction. Dock7 is a RacGEF that has previously been shown to interact with the microtubule cytoskeleton and to influence interkinetic nuclear migration in neural progenitor cells (Watabe-Uchida et al., 2006; Yang et al., 2012). Moreover, like Pcdh19, mutations in human DOCK7 have been linked to an infantile epileptic encephalopathy (Perrault et al., 2014). We cloned zebrafish *dock7* and generated a Dock7-GFP fusion protein. When cotransfected in HEK293, cells Dock7-GFP coimmunoprecipitated Pcdh19-HA (**Figure 3A**). Immunocytochemistry revealed the presence of Pcdh19-HA on the cell surface, as well as in intracellular compartments (**Figure 3C,D**). Dock7-GFP exhibited a cytoplasmic distribution, but showed regions of colocalization with Pcdh19-HA on the cell surface (**Figure 3B,D**).

**Figure 3.**
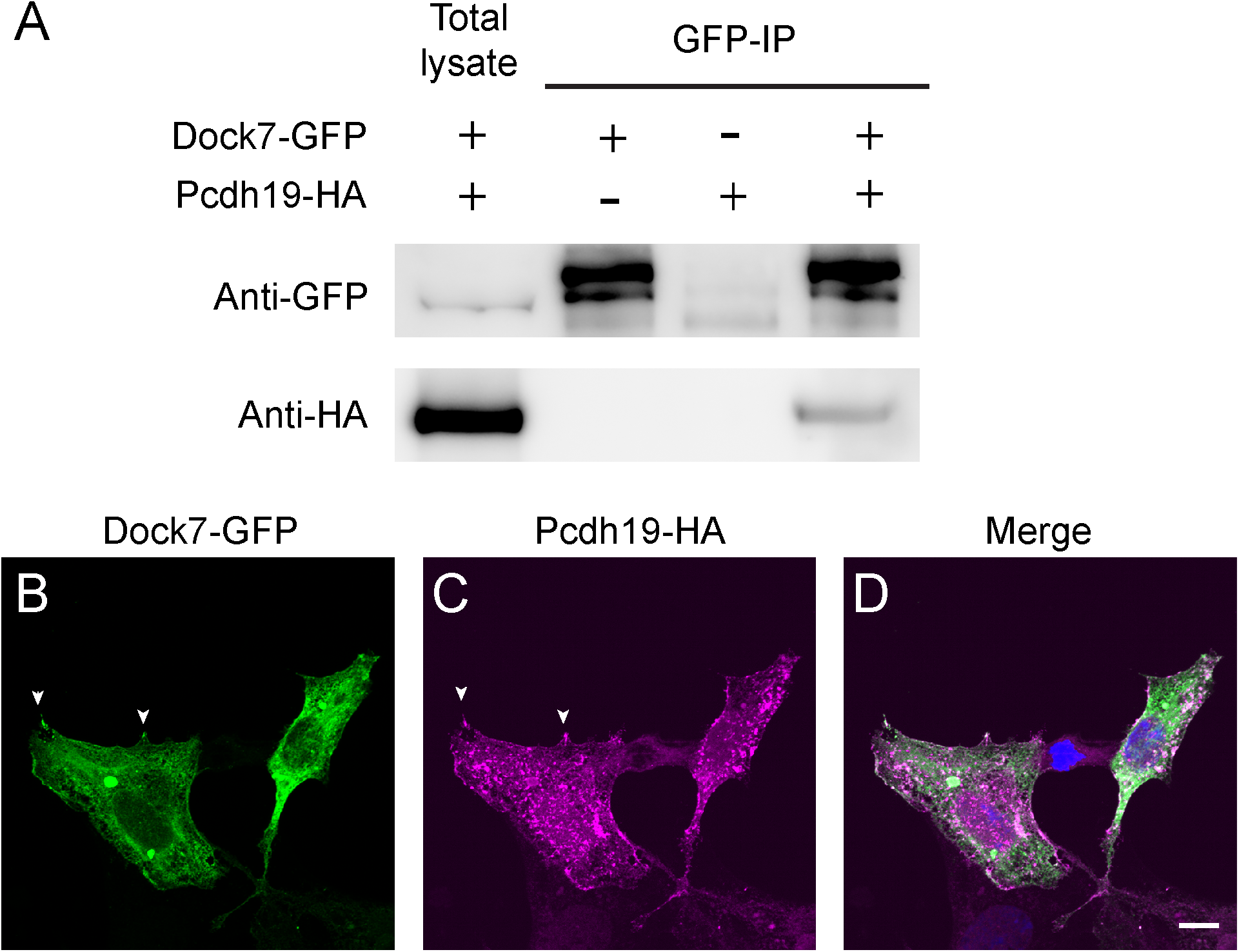
Protocadherin-19 interacts and colocalizes with Dock7. **A**. Zebrafish Dock7-GFP coimmunoprecipitates zebrafish Pcdh19-HA when cotransfected into HEK293 cells. Lysates were incubated with GFP antibody and pulled down using Protein A magnetic beads. Western blots were probed with antibodies against both HA and GFP, confirming an interaction between Pcdh19 and Dock7. **B-D**. Dock7-GFP (**B**,**D**) was cotransfected into HEK293 cells with Pcdh19-HA (**C**,**D**). Though the subcellular patterns of Pcdh19 and Dock7 are distinct, they exhibit some overlap of punctate labeling, particularly at protrusions along the cell surface (arrowheads). Scale bar = *10μ*m.

We previously showed that loss of Pcdh19 results in increased cell proliferation and neuron production in the zebrafish optic tectum (Cooper et al., 2015). Other studies have also indicated a role for Pcdh19 in cell proliferation and differentiation (Fujitani et al., 2016). Consistent with these studies, we observed that Pcdh19 interactors are enriched for proteins associated with microtubules and that are involved with regulating cell divisions. Among these is Neural precursor cell expressed developmentally down-regulated protein 1 (NEDD1), a protein that is associated with the centrosome and is important for spindle function (Pinyol et al., 2013; Yonezawa et al., 2015). In zebrafish, knockdown of *nedd1* leads to increases in both proliferation and apoptosis, resulting in defects in neural tube morphogenesis (Manning et al., 2010). We cloned zebrafish *nedd1* and generated a Nedd1-GFP fusion protein. When cotransfected into HEK293 cells, Pcdh19-HA coimmunopreciptated Nedd1-GFP (**Figure 4A**). To verify the interaction of Pcdh19 with Nedd1 *in vivo*, we expressed Nedd1-HA in zebrafish embryos. Endogenous Pcdh19 co-immunoprecipated with Nedd1-HA from embryo extracts (**Figure 4B**). Immunocytochemistry showed colocalization of Nedd1 and Pcdh19 at cell-cell junctions (**Figure 4C**). Thus, zebrafish Dock7 and Nedd1 can co-exist with Pcdh19 within protein complexes.

**Figure 4.**
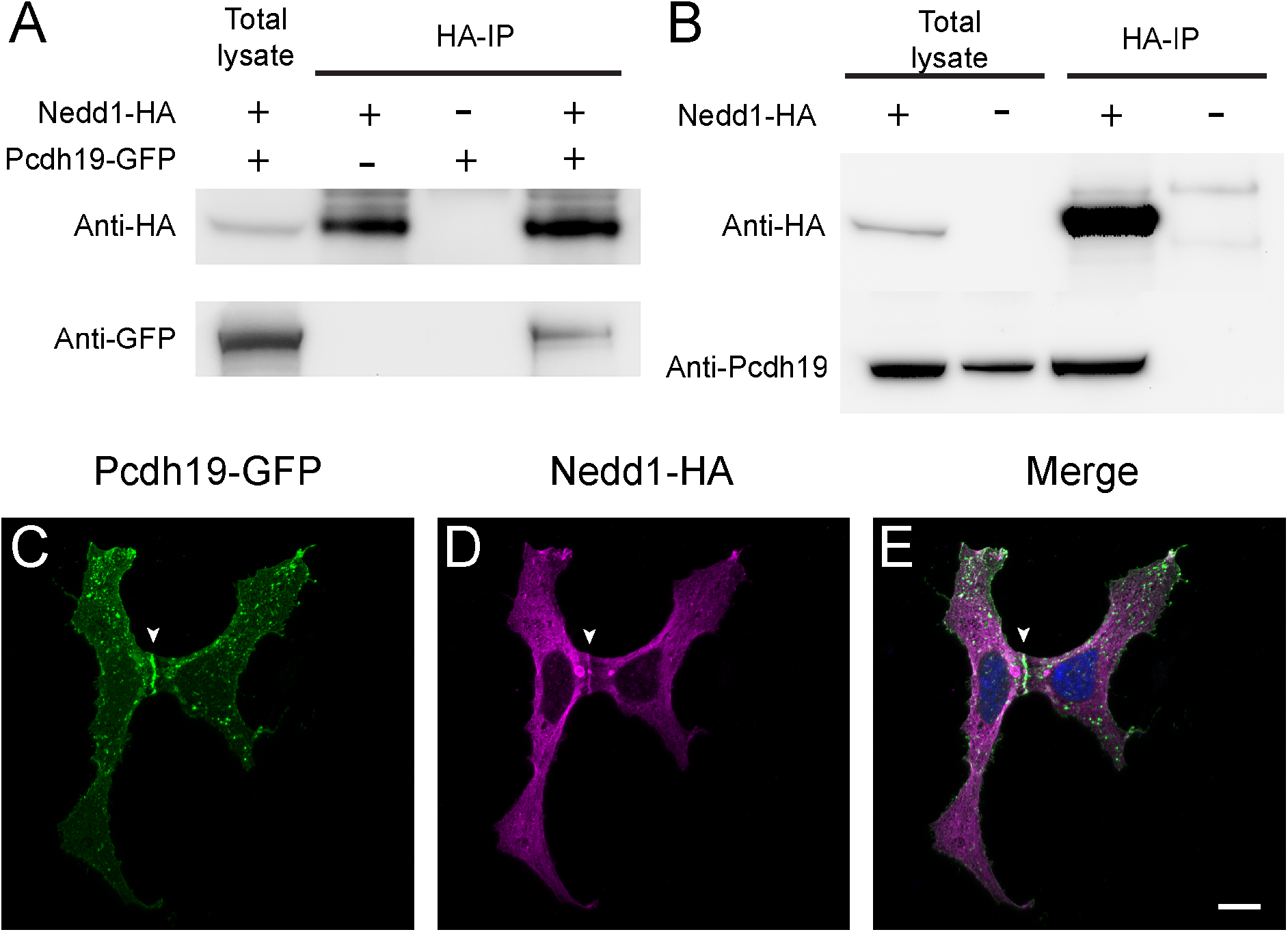
Protocadherin-19 with interacts and colocalizes with NEDD1. **A**. HEK293 cells were cotransfected with zebrafish Nedd1-GFP and zebrafish Pcdh19-HA. Proteins were pulled down from lysates using HA-conjugated magnetic beads and subjected to Western blot analysis. Immunodetection using antibodies against HA and GFP verify an interaction between Pcdh19 and Nedd1. **B**. Nedd1-HA was expressed in zebrafish embryos and extracts were prepared at 24 hours post-fertilization. Nedd1-HA was immunoprecipitated and Western blots were performed using antibodies against HA and endogenous Pcdh19. **C-E**. Pcdh19-GFP(**C**,**E**) was cotransfected into HEK293 cells with NEDD1-HA (**D**,**E**). Though the subcellular patterns of Pcdh19 and NEDD1 are distinct, they exhibit overlap at protrusions along the cell surface, as well as at cell-cell junctions (arrowheads). Scale bar = 10*μ*m.

## DISCUSSION

Mutations in *PCDH19* result in a female-limited infantile epileptic encephalopathy (Depienne et al., 2009; Dibbens et al., 2008), constituting the second most common genetically-based form of epilepsy (Depienne and LeGuern, 2012). Despite progress in understanding the molecular basis of cell adhesion by Pcdh19 and other δ-pcdhs and an advancing understanding in the developmental roles of these proteins, the intracellular pathways downstream of their cell surface interactions are not well defined. Defining the networks of intracellular proteins with which Pcdh19 interacts will both reveal the likely cellular roles of Pcdh19 and provide insight into the molecular mechanisms. This is essential for understanding the biology of Pcdh19, and other δ-pcdhs, and could help elucidate how mutations in *PCDH19* lead to epilepsy. The δ2-pcdhs interact directly with the WAVE complex and promote GTP-Rac1 dependent actin assembly (Biswas et al., 2014; Chen et al., 2014; Hayashi et al., 2014; Nakao et al., 2008; Tai et al., 2010), consistent with a role in contact-dependent cell motility. However, several observations also suggest that the δ-pcdhs regulate the proliferation of neural progenitor cells and the production of neurons (Cooper et al., 2015; Zhang et al., 2014). These results are bolstered by observations implicating the δ-pcdhs as tumor suppressors in a variety of cancers (Xu et al., 2015; Yin et al., 2016; Zong et al., 2017), where forced expression of the δ-pcdhs can inhibit cell proliferation in cancer cell lines. The mechanisms by which δ-pcdhs could regulate proliferation are not known. Here, we used proximity-dependent biotinylation and proteomic analysis to explore the Pcdh19 interactome.

Consistent with the known interactions of the δ2-pcdhs with the WAVE complex, we identified CYFIP1, ABI2 and WASF2 among the Pcdh19 interacting proteins. In addition, we found a number of other actin-associated proteins, such as Afadin, Myosin-VI and Shroom3. In addition to these actin binding proteins, we also identified several regulators of Rho GTPases, which are known to influence actin stability and dynamics. Many of these proteins are regulators of Rac1, consistent with a role for Pcdh19 in controlling Rac1-dependent actin assembly by the WAVE complex. Dock7, in particular, is an intriguing candidate, as it is a RacGEF and mutations in human *DOCK7* have been linked to epilepsy (Perrault et al., 2014). Future functional studies will be required to establish the functional relationship between Dock7 and Pcdh19, and determine their cellular roles during development.

An emerging role for δ2-pcdhs is the regulation of proliferation of neural progenitor cells and subsequent neurogenesis. However, it is not known how adhesion by δ2-pcdhs influences cell division and differentiation. The enrichment of microtubule binding proteins and regulators of cell division suggests that Pcdh19 could play a role in directly regulating the microtubule cytoskeleton in neural progenitor cells. Changes in the adhesive state of δ2-pcdhs could transmit direct and/or indirect signals to the microtubule network. As elements in both the Wnt/β-catenin and Hippo signalling pathways were also identified, the δ2-pcdhs could coordinate both changes in the mechanical state of neural progenitor cells and changes in the signalling pathways that determine cell proliferation and differentiation. The use of BioID only labels proteins within a short distance of the tagged target protein. Thus, this approach does not distinguish between proteins that interact directly from those that are maintained near one another by association in a larger protein complex; future work will be required to identify direct Pcdh19 binding partners. Moreover, further *in vivo* functional analysis will be required in order to fully understand the cellular roles of δ2-pcdhs, including Pcdh19, and their detailed molecular mechanisms.

## METHODS

### Constructs

The coding sequences for BioID-HA and BioID2-HA were obtained from Addgene (www.addgene.org). pcDNA3.1 MCS-BirA(R118G)-HA was a gift from Kyle Roux (Addgene plasmid #36047; http://n2t.net/addgene:36047; RRID:Addgene_36047), and MCS-BioID2-HA was a gift from Kyle Roux (Addgene plasmid #74224; http://n2t.net/addgene:74224; RRID:Addgene_74224). The coding sequence for BioID-HA was amplified using primers containing BamHI and NotI restriction sites and inserted to replace GFP in pEGFP-N1-Pcdh19-GFP (Biswas et al., 2010).

Primers:

BioID-BamHI-F: 5’-gcccgggatccaccggtcaaggacaacaccgtgcccctg-3’;

BioID-NotI-R: 5’-gcggccgcctatgcgtaatccggtacatc-3’;

BioID2-BamHI-F: 5’-gcccgggatccaccggtcgccaccttcaagaacctgatctggc-3’;

BioID2-NotI-R: 5’-gcggccgcctatgcgtaatccggtacatc-3’.

The coding sequences for zebrafish *dock7* and *nedd1* were amplified from cDNA generated from 3 days post fertilization (dpf) embryos (Superscript IV, Thermo Fisher Scientific, Waltham, MA, USA). The sequences were inserted into either the pEGFP-N1 vector (Clontech, Mountain View, CA, USA) or one modified to contain the -HA epitope using primers containing restriction sites.

Primers:

Nedd1-HindII-F: 5’-cgaagcttgccaccatggaggacgtcacacggctg-3’;

Nedd1-BamHI-R: 5’-cgcggatcccctccaccatagttggctcgtagtctttt-3’;

Dock7-NheI-F: 5’-gcggctagcgccaccatggctgagcgccgcgcctttgcc-3’;

Dock7-AgeI-R: 5’-cgcaccggtcctccacctcccatgtctagcttgcgaaggc-3’..

### HEK293 cell culture and calcium phosphate transfection

HEK-293 (ATCC^®^ CRL-1573™,Manassas, VA, USA) cells were maintained in growth media (DMEM with 10% fetal bovine serum and penicillin/streptomycin) at 37°C with 5% CO_2_. Cells were transfected once they reached 70-80% confluence. On the day of transfection, growth media was prewarmed and exchanged 30-60 minutes before calcium phosphate transfection. The DNA/calcium phosphate precipitate was prepared by mixing DNA with 250 mm CaCl_2_ and adding to an equal volume of 2× HBS (280 mm NaCl, 50 mm HEPES, 1.5 mm Na_2_HPO_4_, pH 7.1) using a vortex mixer. The DNA/calcium phosphate mixture was then added dropwise to the cultured cells, which were then incubated at 37°C overnight.

### Immunofluorescence

Transfected HEK293 cells on glass coverslips were rinsed 24 hours following transfection two times in prewarmed phosphate buffered saline (PBS), fresh growth media supplemented with 50 μM biotin was added and the cells incubated overnight at 37°C. The following day, the cells were rinsed two times in PBS, fixed in 4% paraformaldehyde in PBS and permeabilized in 0.25% Triton X-100/PBS for 5 minutes. Cells were then rinsed in PBS and blocked in immunoblock buffer (2% goat serum, 3% bovine serum albumin, 1X PBS) before the addition of antibodies (1:400 mouse anti-HA (Thermo Fisher Scientific) or Streptavidin-HRP (Genscript, Piscataway, NJ, USA)) and incubated overnight at 4°C. Secondary antibody was used to visualize the HA epitope (1:50 goat anti-mouse Alexa 594 (Thermo Fisher Scientific)). To amplify the Streptavidin-HRP signal, tyramide signal amplification was used (TSA-FITC) according to manufacturer’s instructions (PerkinElmer). DAPI (Thermo Fisher Scientific) was added to all coverslips to visualize cell nuclei. Coverslips were rinsed and mounted in Fluoromount G (Electron Microscopy Science, Hatfield, PA, USA) and imaged on a Nikon TiE inverted epifluorescence microscope equipped with a 100x/1.4NA Plan-Apochromat VC oil immersion objective, a Lumencor SOLA LED light source for epifluorescence illumination, and an Andor iXon Ultra 897 EMCCD camera.

### Coimmunopreciptation and Western blotting

HEK293 cells were transiently transfected with cDNAs encoding GFP- or HA-tagged zebrafish Pcdh19, Nedd1 and/or Dock7 using calcium phosphate precipitation as described above. After 24 hours, cells were rinsed in PBS and lysed on ice in cell lysis buffer (20 mM Tris, pH 7.5, 150 mM NaCl, 1 mM EDTA, 0.5% Triton X-100, 1 mM PMSF, and 1X Complete protease inhibitor cocktail (Roche Applied Science, Indianapolis, IN, USA)) and microcentrifuged at 4°C for 10 min. Supernatants were incubated with anti-HA magnetic beads (Thermo Fisher Scientific) for 30-60 minutes at 4°C. For GFP coimmunoprecipitation, the lysates were incubated for 1 hour with 2 μg anti-GFP antibody (Thermo Fisher Scientific) and then incubated at 4°C for 30-60 minutes with Protein A Dynabeads (Thermo Fisher Scientific). The beads were washed five times in wash buffer (20 mM Tris, pH 7.5, 150 mM NaCl, 0.5% Triton X-100), resuspended in loading buffer, and boiled for 5 minutes. Samples were loaded onto 10% Bis-Tris NuPAGE gels (Thermo Fisher Scientific) and subjected to electrophoresis. Proteins were then transferred (Bio-Rad Laboratories, Hercules, CA, USA) to PVDF (GE Life Science, Piscataway, NJ, USA), blocked with 5% nonfat milk in TBST, and incubated overnight with primary antibody (Thermo Fisher rabbit anti-GFP, 1:1,000; Thermo Fisher mouse anti-HA, 1:5,000). HRP-conjugated secondary antibodies (Jackson ImmunoResearch Laboratories, West Grove, PA, USA) were used at 1:5,000, and the chemiluminescent signal was amplified using Western Lightning Ultra (PerkinElmer, Waltham, MA, USA). Blots were imaged on a molecular imaging system (Omega 12iC; UltraLum, Inc., Claremont, CA, USA). For the in vivo co-immunoprecipitation experiments, Nedd1-HA was expressed by co-injecting plasmids *hsp70:Gal4-VP16* and *UAS:Nedd1-HA* into 1-cell stage embryos. Injected embryos were heat shocked at 37°C for 1 hour, beginning at 14 hours post-fertilization (hpf), then embryo extracts were prepared at 18 hpf by homogenizing ∼100 embryos in 500 μL of lysis buffer. Western blots were performed as described above.

### Affinity purification of biotinylated proteins

HEK293 cells (3 × 100 mm plates) were transfected as described above. The cells were rinsed two times in prewarmed PBS 24 hours after transfection and then incubated overnight at 37°C in fresh growth media supplemented with 50 μM biotin. The following day, the cells were rinsed briefly in PBS and then lysed on ice in 1 ml lysis buffer (20 mM Tris, pH 7.5, 150 mM NaCl, 0.5% Triton X-100, 1 mM EDTA, 1 mM PMSF, 1x Complete protease inhibitor (Roche Applied Science)). Cell lysates were recovered following centrifugation and incubated overnight with 50 μL Streptavidin magnetic beads (Cell Signaling Technology, Danvers, MA, USA). Beads were rinsed using a magnetic stand at 25°C according to the following steps: 2 washes for 8 minutes each in 1 ml Buffer 1 (2% SDS in dH_2_0); 1 wash for 8 minutes in Buffer 2 (0.1% deoxycholate, 1% Triton X-100, 500 mM NaCl, 1 mM EDTA, and 50 mM Hepes, pH 7.5); 1 wash for 8 minutes in Buffer 3 (250 mM LiCl, 0.5% NP-40, 0.5% deoxycholate, 1 mM EDTA, and 10 mM Tris, pH 8.1); 2 washes for 8 minutes each in Buffer 4 (50 mM Tris, pH 7.4, and 50 mM NaCl). The magnetic beads were then pooled in 1 ml sterile filtered PBS and analyzed by mass spectroscopy.

### Protein identification using mass spectroscopy

The proteomics analysis was carried out in the OSU Mass Spectrometry and Proteomics Facility in the Campus Chemical Instrumentation Center. The streptavidin magnetic beads were rinsed twice with 50 mM ammonium bicarbonate and the supernatants pooled and reserved. For a third rinse, the beads were left in the supernatant and 5 μL of DTT (5 mg/ml DTT in 50 mM ammonium bicarbonate) was added. The sample was incubated at 56°C for 15 minutes before 5 μL of iodoacetamide (15mg/mL in 50 mM ammonium bicarbonate) was added and incubated for 30 min in the dark at room temperature. Sequencing grade-modified trypsin (500 ng, Promega, Madison, WI, USA) was prepared in 50 mM ammonium bicarbonate and added to the sample, which was allowed to digest overnight at 37°C. Acetic acid was added to quench the enzymatic reaction, the supernatant was concentrated and the peptide concentration was measured using Nanodrop 2000 (Thermo Fisher Scientific).

Capillary-liquid chromatography-nanospray tandem mass spectrometry (Capillary-LC/MS/MS) of global protein identification was performed on a Thermo Fisher Fusion mass spectrometer equipped with a Thermo Easy-Spray operated in positive ion mode. Samples were separated on a Thermo Nano C18 column using an UltiMate 3000 HPLC system from Thermo Fisher Scientific. Each sample was injected into the µ-Precolumn Cartridge (Thermo Fisher Scientific) and desalted with 50 mM acetic acid for 5 minutes. The injector port was then switched to inject and the peptides were eluted off of the trap onto the column. Mobile phase A was 0.1% Formic acid in water and acetonitrile was used as mobile phase B. Flow rate was set at 300 nl/min. Typically, mobile phase B was kept for 2% for 5 minutes before being increased from 2% to 35% in 120 minutes and then increased from 35% to 55% in 23 minutes. Mobile phase B was again increased from 55%-90% in 10 minutes and kept at 90% for another 2 minutes before being brought back quickly to 2% in 2 min. The column was equilibrated at 2% of mobile phase B (or 98% A) for 15 minutes before the next sample injection. MS/MS data was acquired with a spray voltage of 1.7 KV and a capillary temperature of 275 °C was used; S-Lens RF level was set at 60%. The scan sequence of the mass spectrometer was based on the preview mode data dependent TopSpeed method with CID and ETD as fragmentation methods: the analysis was programmed for a full scan recorded between *m/z* 400 – 1600 and a MS/MS scan to generate product ion spectra to determine amino acid sequence in consecutive scans of the five most abundant peaks in the spectrum. To achieve high mass accuracy MS determination, the full scan was performed at FT mode and the resolution was set at 120,000. The AGC Target ion number for FT full scan was set at 4 x 10^5^ ions, maximum ion injection time was set at 50 ms and micro scan number was set at 1. MSn was performed using ion trap mode to ensure the highest signal intensity of MSn spectra. The AGC Target ion number for ion trap MSn scan was set at 100 ions, maximum ion injection time was set at 250 ms and micro scan number was set at 1. The CID fragmentation energy was set to 35%. Dynamic exclusion is enabled with a repeat count of 1, exclusion duration of 60s and a low mass width and high mass width of 10ppm.

Sequence information from the MS/MS data was processed by converting the .raw files into a merged file (.mgf) using MSConvert (ProteoWizard). The resulting mgf files were searched using Mascot Daemon by Matrix Science version 2.5.1 (Boston, MA) and the database searched against the most recent databases. The mass accuracy of the precursor ions was set to 10ppm, accidental pick of 1 ^13^C peaks was also included into the search. The fragment mass tolerance was set to 0.8 Da. Considered variable modifications were oxidation (Met), deamidation (N and Q) and carbamidomethylation (Cys). Four missed cleavages for the enzyme were permitted. A decoy database was also searched to determine the false discovery rate (FDR) and peptides were filtered according to the FDR. The significance threshold was set at p<0.05 and bold red peptides was required for valid peptide identification. Proteins with a Mascot score of 50 or higher with a minimum of two unique peptides from one protein having a *-b* or *-y* ion sequence tag of five residues or better were accepted. Any modifications or low score peptide/protein identifications were manually checked for validation.

Two experiments were performed with Pcdh19-BioID-HA and three experiments were performed with Pcdh19-BioID2-HA. We manually removed keratins, ribosomal proteins and nuclear proteins from each datset. In addition, we used the Crapome database (www.crapome.org) to remove proteins that were present in more than 15% of reported proteomics datasets. Finally, proteins that were identified in 3 out of the 5 experiments were retained in a final list of Pcdh19 interacting proteins. Gene ontology analysis was performed with the Cytoscape (https://cytoscape.org) plugin ClueGO (Bindea et al., 2009). For analysis, we used GO term fusion and pathways with p<0.05. We additionally submitted our protein list to the String database (www.string-db.org).

## Acknowledgments

This work was supported by the National Science Foundation Grant IOS 1457126, an NIH Grant R01 EY027003, NIH Shared Instrumentation Grants S10 OD010383, S10 OD18056, and an NIH Neurosciences Center Core Grant P30 NS104177.

